# Measuring the Landscape of CpG Methylation of Individual Repetitive Elements

**DOI:** 10.1101/018531

**Authors:** Yuta Suzuki, Jonas Korlach, Stephen W. Turner, Tatsuya Tsukahara, Junko Taniguchi, Wei Qu, Kazuki Ichikawa, Jun Yoshimura, Hideaki Yurino, Yuji Takahashi, Jun Mitsui, Hiroyuki Ishiura, Shoji Tsuji, Hiroyuki Takeda, Shinichi Morishita

## Abstract

Determining the methylation state of regions with high copy numbers is challenging for second-generation sequencing, because the read length is insufficient to map reads uniquely, especially when repetitive regions are long and nearly identical to each other. Single-molecule real-time (SMRT) sequencing is a promising method for observing such regions, because it is not vulnerable to GC bias, it performs long read lengths, and its kinetic information is sensitive to DNA modifications. We propose a novel algorithm that combines the kinetic information for neighboring CpG sites and increases the confidence in identifying the methylation states of those sites. Both the sensitivity and precision of our algorithm were ∼93.7% on CpG site basis for the genome of an inbred medaka (*Oryzias latipes*) strain within a practical read coverage of ∼30-fold. The method is quantitatively accurate because we observed a high correlation coefficient (*R* = 0.884) between our method and bisulfite sequencing, and 92.0% of CpG sites were in concordance within 0.25. Using this method, we characterized the landscape of the methylation status of repetitive elements, such as LINEs, in the human genome, thereby revealing the strong correlation between CpG density and unmethylation and detecting unmethylation hot spots of LTRs and LINEs. We could uncover the methylation states for nearly identical active transposons, two novel LINE insertions of identity ∼99% and length 6050 base pairs (bp) in the human genome, and sixteen *Tol2* elements of identity >99.8% and length 4682 bp in the medaka genome.

## 1 BACKGROUNDS

There has been a great deal of interest in identification of genome-wide epigenetic DNA modifications in recent years, because DNA modifications play an essential role in cellular and developmental processes (Weaver et al., 2004; Anway et al., 2005; Jirtle and Skinner, 2007; Miller, 2010; Zemach et al., 2010; Schmitz et al., 2011; Molaro et al., 2011; Smith et al., 2012; Qu et al., 2012). Some of human transposable elements, such as long interspersed nuclear elements (LINE), are reported to transpose actively within somatic cells along differentiation of neural tissues, and to be partly regulated by DNA methylation (Muotri et al., 2005, 2010). Each family of human transposable elements is in a variety of methylation statuses according to tissue type by looking at the mixture of methylation information on the consensus sequence of TEs in the same family (Xie et al., 2013). Many human diseases are also associated with DNA methylation states of transposable elements. In particular, unmethylation of repetitive elements, such as LINE-1 elements, has been related to some cancers (Wilson et al., 2007; Ross et al., 2010). Although only a few LINE-1 elements exhibit activity in the human genome (Beck et al., 2010), transpositions of these elements have been reported in various cancer genomes (Lee et al., 2012; Goodier, 2014), and importantly, it has been reported that transpositions are correlated with unmethylation of the promoter region of LINE-1 elements (Tubio et al., 2014). Therefore, it is essential to develop an experimental framework that can characterize the methylation state of repetitive elements in a genome-wide manner.

The advent of second-generation sequencing technology has increased the efficiency of the generation of precise genome-wide methylation maps at a single-base resolution using bisulfite treatment (Cokus et al., 2008; Lister et al., 2008; Meissner et al., 2008; Lister et al., 2009; Harris et al., 2010); however, these sequencing-based technologies have difficulty in characterizing the methylation status of CpGs in regions that are highly similar to other regions. Bisulfite-treated short reads from these regions often fail to map uniquely to their original positions; instead, they are likely to be aligned ambiguously to multiple genomic positions. Moreover, first/second-generation sequencing technology often fails to sequence DNA regions with a GC content >60% (Aird et al., 2011; Ross et al., 2013) and may exhibit bias against GC-rich regions. These inherent problems of second-generation sequencing may result in underrepresentation of methylation information on specific DNA regions, such as transposable elements and low-complexity repeat sequences (Lister et al., 2009; Harris et al., 2010; Bock et al., 2010; Gifford et al., 2013; Jiang et al., 2013). Especially, the younger and more active transposons are thought to retain higher fidelity and are therefore difficult to address using short reads.

In the PacBio RS sequencing system, DNA polymerase is used to perform single-molecule real-time (SMRT) sequencing (Korlach et al., 2008; Eid et al., 2009), and this system is capable of sequencing reads of an average length of >10 kb. SMRT sequencing is also able to sequence genomic regions with extremely high GC contents. A striking example is a previous report of the sequencing of a >2-kb region with a GC content of 100% (Loomis et al., 2012), indicating that SMRT sequencing is less vulnerable to sequence composition bias than is first/second-generation sequencing. SMRT sequencing of bisulfite-treated DNA fragments may allow identification of DNA methylation within long regions; however, this approach is not promising because bisulfite treatment divides DNA into short fragments <1000 bp (Miura et al., 2012).

Instead, we explored another advantage of SMRT sequencing to detect DNA modifications. In SMRT sequencing, we can observe the base sequence in a single DNA molecule as each corresponding nucleotide is incorporated using the time course of the fluorescence pulses. From this time course information, we can determine the inter-pulse duration (IPD), which is defined as the time interval separating the pulses of two neighboring bases. Importantly, the IPD of the same genomic position varies and has a significant and predictable response to DNA modifications due to the sensitivity of DNA polymerase kinetics to DNA modifications and damage (Flusberg et al., 2010).

Consequently, the IPD ratio (IPDR), the ratio of the average IPD in DNA templates with modifications to that in control templates, tends to be perturbed systematically, allowing identification of DNA modifications (Fig. 1A). Indeed, SMRT sequencing methods have been used to detect changes in 5-hydroxymethylcytosine (Flusberg et al., 2010), N4-methylcytosine (Clark et al., 2012), and N6-methylademine (Flusberg et al., 2010; Fang et al., 2012; Feng et al., 2013; Greer et al., 2015), as well as damaged DNA bases (Clark et al., 2011) in bacteria and mitochondria; however, estimation of 5-methylcytosine (5-mC) residues using low-coverage reads is prone to errors and requires extensive coverage at each position to clarify the base-wise 5-mC state and therefore becomes costly (Flusberg et al., 2010; Fang et al., 2012; Schadt et al., 2012). Clark *et al.* attempted to improve the detection of microbial 5-mC in the *Escherichia coli* and *Bacillus halodurans* genomes using Tet1-mediated oxidation to convert 5-mC into 5caC in SMRT reads of ∼150x coverage per DNA strand (Clark et al., 2013). Therefore, kinetic information from low-coverage SMRT reads at a single CpG site is not reliable for predicting the methylation status.

**Figure 1.**
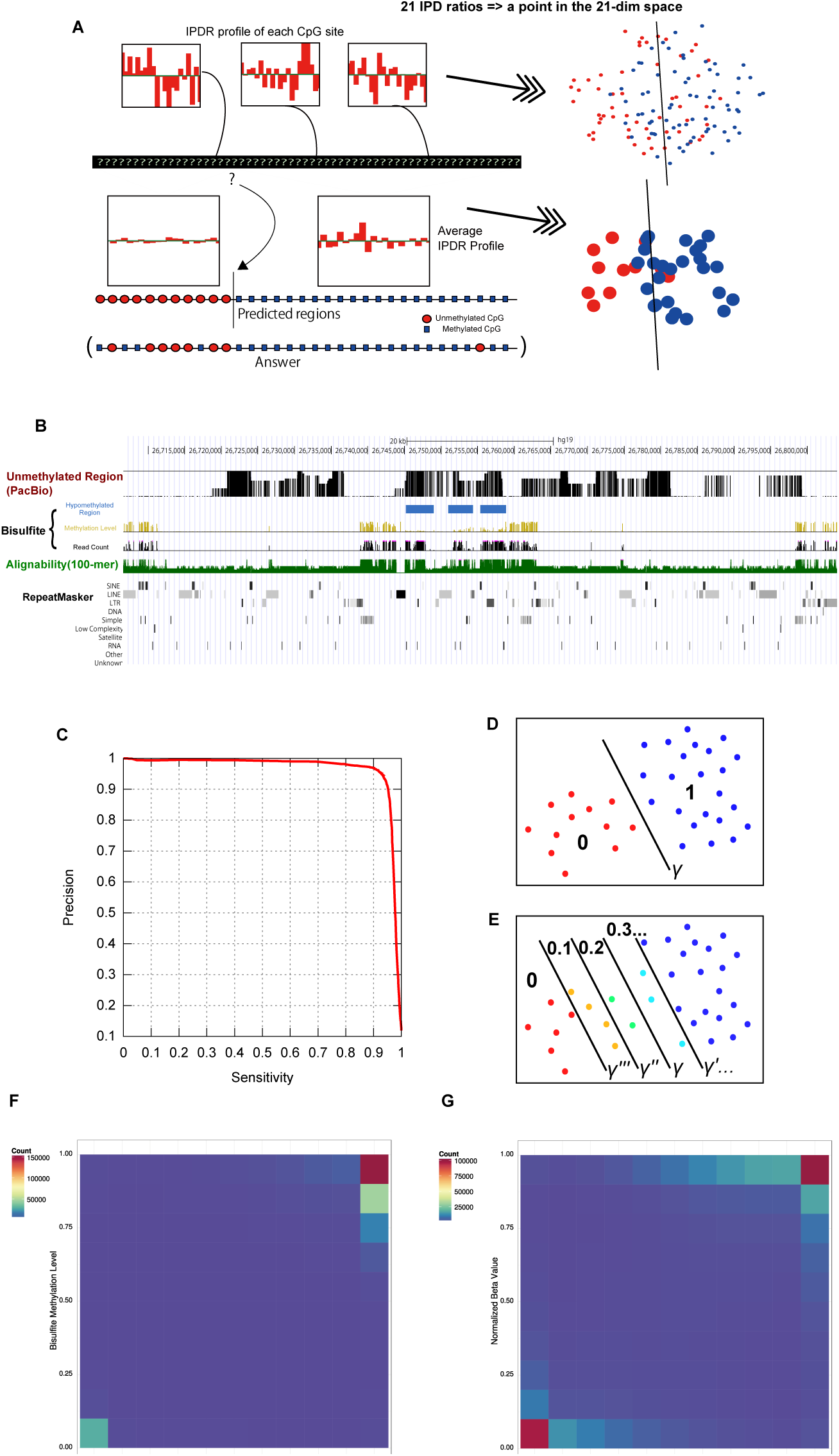
Outline of our integration method. **A.** The top three distributions show the typical Inter-Pulse Duration Ratio (IPDR) profiles within 10 bp of the CpG sites in the raw data. The IPDR profiles of individual CpG sites were treated as points in the 21-dimensional feature space. Red-colored unmethylated CpGs and blue-colored methylated CpGs are difficult to separate using a hyperplane. Therefore, initially, we had little knowledge about the methylation status of each CpG site from the raw data, as illustrated by the question marks at the CpG sites. Our algorithm predicts the boundary of unmethylated and methylated CpG sites. The average IPDR profiles of the two regions, which have clearly distinct IPDR profiles, are shown below the two regions separated by the boundary (see the detailed IPDR profiles in Supplemental Fig. S1B). Red circles and blue boxes represent unmethylated and methylated CpGs, respectively, predicted by our algorithm (annotated as ‘predicted regions’) and were observed by bisulfite sequencing (labeled ‘answer’). In the feature space, red and blue disks represent the IPDR profiles of predicted regions. **B.** Comparison of our prediction with the available human genome methylome data. From top to bottom, black bars indicate unmethylated regions predicted from SMRT sequencing data using our method. Yellow and black bars show the methylation level and read coverage obtained from public bisulfite sequencing data, respectively, and blue boxes show unmethylated regions predicted from the bisulfite data. Green bars below indicate the alignability of short (100-bp) reads. The bottom row shows repeat masker tracks. Both methods are consistent in showing unmethylation on the three blue-colored regions. No read counts of the bisulfite data are available in long duplicated regions where the alignability is quite low, but our method can estimate unmethylation in these regions. **C.** The sensitivity and precision (proportion of true-positives among the predicted positives) of our integration method are evaluated on individual CpG sites when we change the intercept of the hyperplane and set the minimum number of CpG sites in a region, *b*, to 35. See the curves for *b* = 30,40,45, and 50 in Supplemental Fig S3. **D.** IPDR profiles of CpGs are represented as points in the feature space. Predictions are made using a decision hyperplane determined by its intercept *γ*, and individual CpGs are classified as methylated (blue) or unmethylated (red). **E.** Multiple predictions using a set of different intercept parameter values define the discrete methylation level (DML) on each CpG site. Specifically, after decomposing DNA into unmethylated and methylated regions for different intercept values of *γ*, we compute the ratio of methylated regions that cover each CpG site, and treat the ratio as the methylation level of the CpG site. **F.** DML (x-axis) and methylation level monitored by bisulfite sequencing (y-axis) in our medaka sample. The colors are based on the log of the number of CpG sites having corresponding DML value and bisulfite methylation level. These values were strongly correlated (*R* = 0.884) and the difference was within 0.25 for 92.0% of CpG sites. Most of the CpG sites were methylated because we observed CpG methylation in a genome-wide manner. **G.** DML (x-axis) correlated (*R* = 0.732) with the normalized beta values of BeadChip (y-axis) for the CpG sites in our human sample, and 75.4% of CpG sites are in concordance within 0.25. The majority of CpG sites are unmethylated, because most CpG sites on the BeadChip are designed on CpG islands.

In this study, we exploited the facts that unmethylated CpG dinucleotides are rare (∼10%) in vertebrates and generally do not exist in isolation but often range over long unmethylated regions (Qu et al., 2012; Gifford et al., 2013; Eckhardt et al., 2006; Bock et al., 2008; Shoemaker et al., 2010; Nautiyal et al., 2010; Xie et al., 2013). Su *et al.* reported that the average length of unmethylated regions in five human cell types is ∼2 kb (Su et al., 2012b). Thus, estimating regions of unmethylated CpG sites is informative in most cases. Similarly, integrating kinetic information for many CpG sites in a long region can increase the confidence in detecting methylation when the status of those sites is correlated and shows promise for predicting the methylation status in a block using low-coverage SMRT reads. We examined the feasibility of the approach and present a novel computational algorithm that integrates SMRT sequencing kinetic data and determines the methylation statuses of CpG sites.

To demonstrate a possible application of our method, we investigated methylation status of individual occurrences of transposable elements in the human genome. Only a few studies have addressed this. In one study, the authors designed a microarray specifically to characterize changes in transposon methylation of cancer cells (Szpakowski et al., 2009), and other study utilized a bisulfite sequencing dataset (Su et al., 2012a). These methods, however, cannot observe the repetitive elements which have not diverged enough to be distinguished by short reads, or those which are novel insertions and simply absent from the reference genome. Therefore, we examined the possibility of determining the methylation statuses of highly similar occurrences of transposable elements in human and medaka fish (*Oryzias latipes*), which could be investigated only by using long reads.

## 2 RESULTS

### 2.1 Bisulfite data benchmark and SMRT sequencing

It is necessary to take into account allele-specific DNA methylation in the analysis of the methylomes of diploid genomes (Shoemaker et al., 2010; Chandler et al., 1987; Yamada et al., 2004; Zhang et al., 2009; Kerkel et al., 2008; Schilling et al., 2009; Hellman and Chess, 2010), because we may observe an intermediate DNA methylation level resulting from the mixture of different methylation states from two haplotypes (Lister et al., 2009; Deng et al., 2009). To assess the ability of SMRT sequencing to monitor the DNA methylation status, DNA extracted from a haploid cell line would serve as an ideal template, avoiding situations in which two alleles are differentially methylated. In nonhuman model organisms, inbred strains also provide a clean resource, because the two haplotypes are almost identical in sequence, suggesting that the methylation statuses of the two haplotypes may also match. Therefore, we used the medaka model system (Kasahara et al., 2007), because six medaka methylomes are available from early embryos, testes, and liver in two inbred strains (Qu et al., 2012) by way of Illumina bisulfite sequencing, which outperforms three other frequently used sequence-based methods in terms of the genome-wide percentage of CpGs covered (Harris et al., 2010). As CpG methylation status reference data, we used the testes methylome of the medaka Hd-rR inbred strain. In this dataset, most of the CpG sites in the medaka genome are either unmethylated or methylated, and methylation at non-CpG sites is very rare (∼0.02%) (Qu et al., 2012), allowing us to focus on CpG sites only. We collected 31.06-fold coverage SMRT subreads from the testes of medaka Hd-rR (assuming an estimated genome size of 800 Mb) using P6-C4 reagents. We also collected 22.45-fold and 13.06-fold coverage SMRT reads from human peripheral blood of two Japanese individuals. Thus, we have 3 datasets in total, 1 for medaka and 2 for human. For sequencing two human samples, we employed the P6-C4 reagents and the P4-C2 or C2-C2 reagents, respectively. In total 2848641, 7279594, and 19115712 subreads were anchored to the medaka genome and the human genome, respectively. The mean mapped subread lengths were 8722 bases for medaka and 9254 and 2049 bases for 2 human samples (Supplemental Table 1).

### 2.2 Prediction of the methylation state from kinetic data

Figure 1A shows a schematic representation of the basic concept of our method. First, as a raw ingredient for prediction, we defined the IPDR profile of a CpG site as an array of IPDR measurements of 21 bp surrounding the CpG site. With low coverage, the IPDR profiles at individual CpG sites are noisy and insufficient for determining whether the focal CpG site is unmethylated or methylated. However, if we could somehow identify the boundaries of unmethylated/methylated regions, it would be possible to take the average of the IPDR profile for the CpGs within each region and would allow better prediction of the methylation state of each region from its average IPDR profile, which has less noise than the profile of a single CpG site.

We implemented our method using linear discrimination of the vectors of (average) IPDR profiles around the focal CpG sites. We represented the vectors as points residing in the Euclidean space of the appropriate dimension and attempted to separate the points by a decision hyperplane. For better accuracy, we optimized two parameters of the decision hyperplane, the orientation and intercept. Supplemental Fig. S1A shows the optimized orientation. As unmethylated regions are ∼2 kb in size (on average) and contain ∼50 CpG sites in vertebrate genomes (Su et al., 2012b), in the prediction, we first assume unmethylated regions to have at least 50 CpG sites and integrate the IPDR profiles to make predictions, which is effective in reducing noise in the IPD measurements. We will later examine whether we can reduce the number of CpG sites, denoted by *b*, while preserving the accuracy. Our method divides the genome into regions containing ≥ *b* CpG sites, such that each region is either unmethylated or methylated. An example of our prediction for the human genome is shown in Fig. 1B, in which our method is able to estimate unmethylation of long duplicated regions while the bisulfite sequencing provides little information. Supplemental Fig. S1C illustrates another example in which both methods are consistent in showing unmethylation in gene promoters.

While setting lower bound *b* to 50 is supported by the plausible heuristics with biological grounds, a looser bound (*b* < 50) allows us to detect shorter regions. We therefore examined whether we could use a smaller value of *b* (= 30, 35, 40, 45) without degrading the accuracy of prediction. To evaluate the accuracy of our method, we used the chromosome 1 of length 34,959,811 bp in the medaka genome (version 2) that we assembled from SMRT subreads. For predicting CpG methylation accurately, we guaranteed that each CpG site was covered by at least three subreads, which slightly reduced the original average read coverage, 31.06-fold, to 29.9-fold on the chromosome 1. To examine the coverage effect, we used five subread sets of coverage 20%, 40%, 60%, 80%, and 100% of 29.9-fold. We calculated various accuracy measures, such as sensitivity (recall), specificity (1−false-positive rate), and precision by comparing our prediction on each CpG site with the methylation level determined in a bisulfite sequencing study (Qu et al., 2012). As most CpG sites in the medaka genome are methylated consistently, there are only a small number of positive examples of unmethylated CpGs, and therefore, precision is more informative than specificity in evaluation. We made the trade-off between sensitivity and precision through the selection of the intercept of the decision hyperplane (Supplemental Fig. S2-3). When we used 100% of 29.9-fold subreads, setting *b* to 35 outperformed the other values (Supplemental Fig. S3), and for better readability, Figure 1C illustrates the sensitivity and precision curve for *b* = 35 only. Our prediction achieved 93.7% sensitivity and 93.9% precision, or 93.0% sensitivity and 94.9% precision, depending on the selection of the intercept. In contrast, for a smaller coverage, a wider window (a larger value of *b*) was favored; for example, for coverages of 20% and 40% of 29.9-fold, setting *b* to 50 performed best (Supplemental Fig. S3). Both sensitivity and precision were ∼ 90% for *b* = 45 even if the coverage is relatively small, 60% of 29.9-fold (Supplemental Fig. S3C). Overall, sensitivity and precision of our method are substantially high using a reasonable coverage of SMRT subreads.

### 2.3 Handling intermediate or ambiguous methylation states

We have introduced a two-class model of our prediction that assigns all of the CpG sites into either unmethylated or methylated regions; however, such a dichotomous model is rather unrealistic, and more refined predictions involving multilevel methylation states or even continuous methylation levels are desirable. For example, an intermediate level of CpG methylation could result from the distinct methylation states of two DNA molecules of diploid cells, although each cytosine must be either methylated or unmethylated in a single DNA molecule. More generally, a sampled cell population can be epigenetically heterogeneous, which would possibly show a spectrum of methylation levels according to its composition. Finally, prediction allowing intermediate states can represent the ambiguity of the prediction, and exclusion of such ambiguous predictions is expected to improve the overall prediction accuracy.

Taking these points into consideration, we extended our method to achieve more complex and informative multi-class predictions. Figures 1D-E depict the concept for multi-class prediction using hypothetical data points. We made a classification using the linear discrimination process involving a separation (decision) hyperplane and determined the position of the hyperplane using the intercept parameter denoted by *γ* (Fig. 1D). Intuitively, the intermediately methylated CpGs are expected to be distributed more closely to the decision plane, and are therefore more ambiguous than CpGs with *bona fide* methylation states are. Thus, to output the multi-class prediction, we perturbed the intercept *γ* around its optimal value to produce multiple predictions on each CpG site, which is illustrated by the parallel displaced hyperplanes (Fig. 1E). We then defined the discrete methylation level (DML) as the fraction of predictions that favored ‘methylation’. The robust predictions on the *bona fide* methylation states should have extreme DML values, unlike intermediate or ambiguous predictions.

We checked the accordance between our DML and intermediate or ambiguous methylation level captured by two other quantitative methods, bisulfite sequencing and Illumina BeadChip. On the medaka sample, we observed a strong correlation (*R* = 0.884) between our DML and methylation level calculated from bisulfite sequencing, and we confirmed that 92.0% of CpG sites were in concordance within 0.25 (Fig. 1F). We also compared our DML on the human sample to the beta value (an indicator of methylation level expressed as a value ranging over [0,1]) obtained from Illumina BeadChip (Fig. 1G). We observed a weaker correlation (*R* = 0.816) and a smaller fraction (75.4%) of CpG sites in concordance within 0.25 presumably because the beta value is less quantitative than the methylation level calculated from bisulfite sequencing (Wang et al., 2015). Overall, DML serves to reflect the quantitative nature of methylation status in the samples.

### 2.4 Genome-wide methylation pattern of repetitive elements in the human genome

We investigated how individual occurrences of repetitive elements were methylated in the human genome, as summarized in Table 1. Of note, some occurrences of repetitive elements contain no or very few CpG sites, and thus we only consider those occurrences with at least 10 CpGs to exclude other less informative cases. First, we checked whether SMRT reads could address the repetitive regions in a useful manner for methylation analysis. Specifically, we considered a repeat occurrence to be covered by uniquely mapped SMRT reads if the IPD ratio was available on ≥50% of CpGs, and found that >96% were covered for every repeat type. To draw robust conclusions, we further applied a stringent quality control process to each repeat occurrence such that the read coverage was >5. Although this step reduced the number of repeat occurrences under consideration by 3−18%, this reduction could be mitigated simply by producing more data. Finally, we treated an occurrence as unmethylated if ≥50% of CpGs were predicted as unmethylated. Fractions of unmethylated repeat occurrences vary considerably among different classes of repetitive elements, from ∼1% for L1 and Alu to ∼50% for MIR and >70% for simple repeats and low-complexity regions. To validate our prediction regarding the repeat occurrences, we selected 21 regions for bisulfite Sanger sequencing, designed primers for nested PCR (Supplemental Table S2), and could amplify six regions, indicating the difficulty in observing DNA methylation of repetitive elements using traditional bisulfite Sanger sequencing. In the six amplified regions, we confirmed the consistency between our prediction and the methylation state observed by bisulfite Sanger sequencing (Supplemental Fig. S4).

We then examined the features for characterizing the differences between methylated and unmethylated repetitive elements. First, CpG density was significantly higher in the unmethylated occurrences in almost all classes of repetitive elements (*p* < 1%, Fig. 2A). This observation was consistent with the known association between CpG-rich regions and unmethylation because methylation leads to depletion of CpG sites through deamination (Cooper and Krawczak, 1989). Second, sequence divergence from the representative in each repeat class also showed a correlation with methylation status (Fig. 2B). For most classes, with the apparent exception of simple repeats, low-complexity regions, and MIR elements, unmethylated occurrences were significantly more divergent than were methylated occurrences (*p* < 1%, Fig. 2B), presumably because younger copies of a repeat element are less divergent and are likely to be targets of DNA methylation. We also examined whether the methylation status of repetitive elements could be correlated partly with sequence features. Kernel principal component analysis (PCA) using spectrum kernel suggested positive answers for some repeat types (Supplemental Fig. S5).

**Figure 2.**
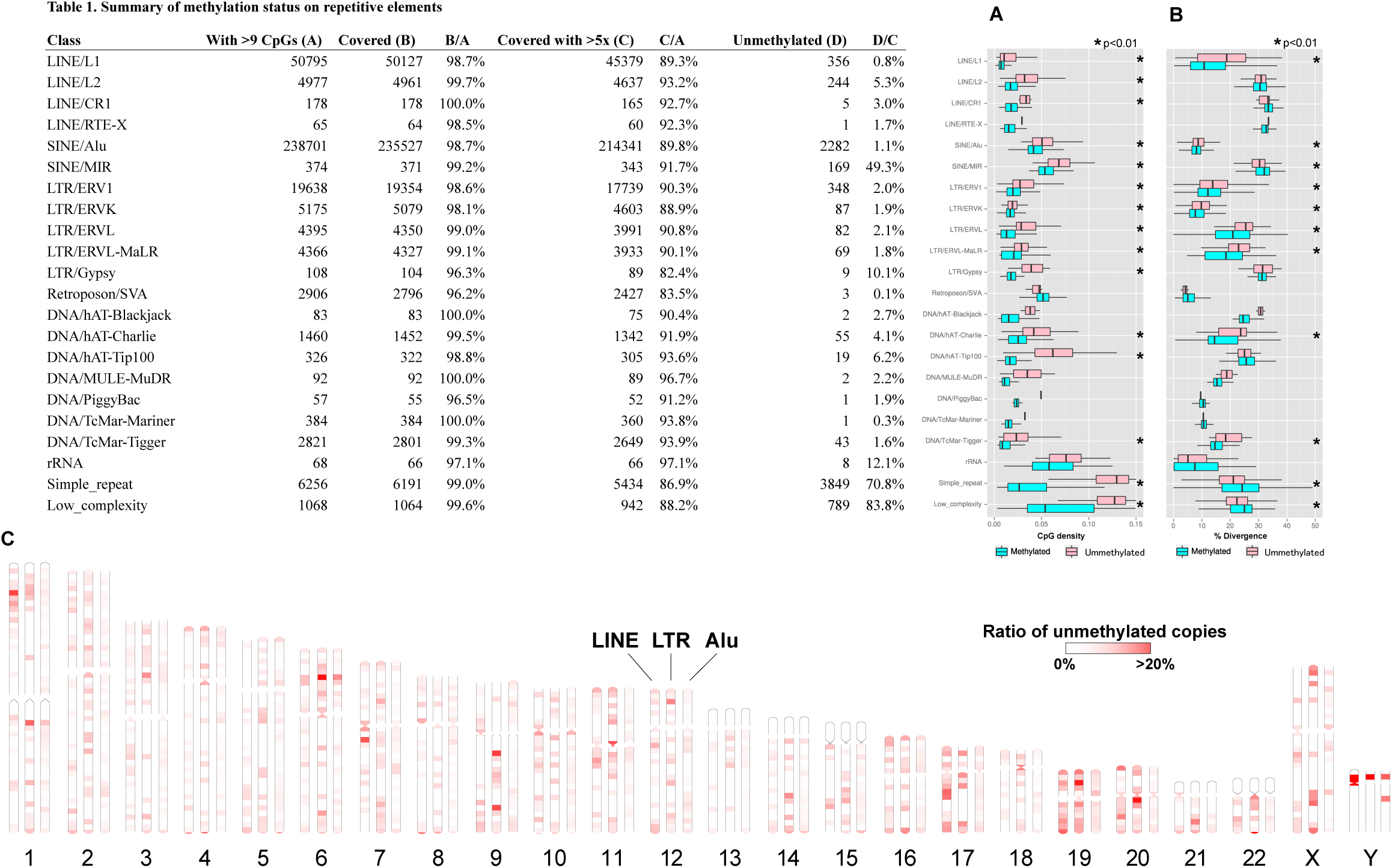
Epigenetic landscape of repetitive elements in the human genome. **A-B.** Distribution of CpG density (**a**) and sequence divergence from the representative in each repeat class (**b**) for methylated (cyan) and unmethylated (pink) repeat occurrences. The asterisks indicate statistical significance (*p* < 1%) determined by the U test. **C.** Genome-wide distribution of unmethylated repetitive elements. The ratio of unmethylated repeat occurrences to all occurrences in each 5-Mb bin is indicated by color shadings. Prediction of the methylation state was performed after quality control as described in the text.

Next, we examined whether the unmethylated repeat occurrences were distributed uniformly or non-uniformly throughout the entire genome. We selected three major classes (LINE, Alu, and LTR) of repetitive elements for this analysis. We calculated the ratios of unmethylated copies to all repetitive elements in individual non-overlapping bins 5 Mb in size (Fig. 2C). The non-random distribution patterns were more evident for LINE and LTR than for Alu. For example, we found unmethylated LINEs to be enriched in the p-arm of chromosome 1 and in chromosomes 17 and 19. There were unmethylation ‘hot spots’ of LTR elements, *e.g.*, in chromosomes 6 and 9 (Supplemental Fig. S6). It is intriguing that some of these unmethylation hot spots, such as those in the p-arms of chromosomes 6 and Y, seem to be shared among different classes of repetitive elements.

We further investigated the methylation states of LINE/L1 elements, the only active autonomous retrotransposons in mammals (Furano, 2000). Although most of LINE/L1 insertions contain many mutations, Penzkofer *et al.* categorize full-length L1 elements into three classes according to the conservation of two open reading frames (ORFs) (Penzkofer et al., 2005); namely, L1s with intact in the two ORFs that are likely to exhibit retro-transposition activity, L1s with an intact ORF2 but a disrupted ORF1, and non-intact L1s. We obtained the positions of these human LINE/L1 elements from L1Base (Penzkofer et al., 2005) and analyzed their methylation states (Supplemental Table 3). Although 0.61% of non-intact L1s were unmethylated, all of L1s with intact in two ORFs and L1s with an intact ORF2 were methylated. We also checked the presence of LINE insertions that were novel to the hg19 reference genome. We assembled the SMRT reads using FALCON and searched the assembly for novel LINE insertions that matched a hot L1 element (GenBank: M80343.1) of size 6050 bp with identity > 98.5%. The hot L1 element was used as the representative according to the procedure of L1Base (Penzkofer et al., 2005). We identified two novel instances covered by sufficient depth of SMRT reads that allowed us to call their methylation statuses confidently. Both of the two LINE insertions were estimated to be methylated, and the positions of these two occurrences are found in Supplemental Fig. S7. These results confirmed putatively active LINE/L1 elements with intact ORFs were targets of DNA methylation.

### 2.5 Analysis of the *Tol2* transposable element in medaka

The medaka has an innate autonomous transposon known as *Tol2*, which is one of the first examples of autonomous transposons in vertebrate genomes and a useful tool for genetic engineering of vertebrates, such as zebrafish and mice (Kawakami, 2007). The excision activities of *Tol2* are promoted when DNA methylation is reduced by 5-azacytidine treatment, which suggests that DNA methylation is one of the mechanisms regulating the *Tol2* transposition (Iida et al., 2006). Nevertheless, observing the methylation status of each *Tol2* copy using short reads is difficult, because *Tol2* is 4682 b in length, and ∼20 highly similar copies of *Tol2* exist in the genome (Koga et al., 2000).

To elucidate the methylation status of each *Tol2* copy, we applied our method to a new assembly of the Hd-rR genome obtained exclusively from SMRT reads. We found 17 copies of *Tol2* contained entirely within this assembly, all of which were essentially identical (>99.8% sequence identity). We then called the methylation status of these *Tol2*. For comparison, we mapped bisulfite-treated short reads to these contigs and determined the methylation level. The methylation status of these *Tol2*, observed by SMRT reads and bisulfite-sequencing, are shown in Fig. 3. While virtually no *Tol2* copies were mapped by bisulfite reads, as expected from their extremely high fidelity, 16 of 17 copies were anchored by SMRT reads, and all were predicted to be methylated by our method. For the regions examined by both SMRT reads and bisulfite-treated short reads, our prediction was consistent with the methylation level calculated from the bisulfite-treated reads. For example, one *Tol2* copy was surrounded by unmethylated regions (number 14). From the bisulfite data, it appeared that the body of *Tol2*, from which data were missing, was unmethylated. Nevertheless, our prediction estimated this region to be methylated. These results demonstrate the ability of our method using SMRT reads to clarify DNA methylation states of highly identical repetitive elements such as active transposons.

**Figure 3.**
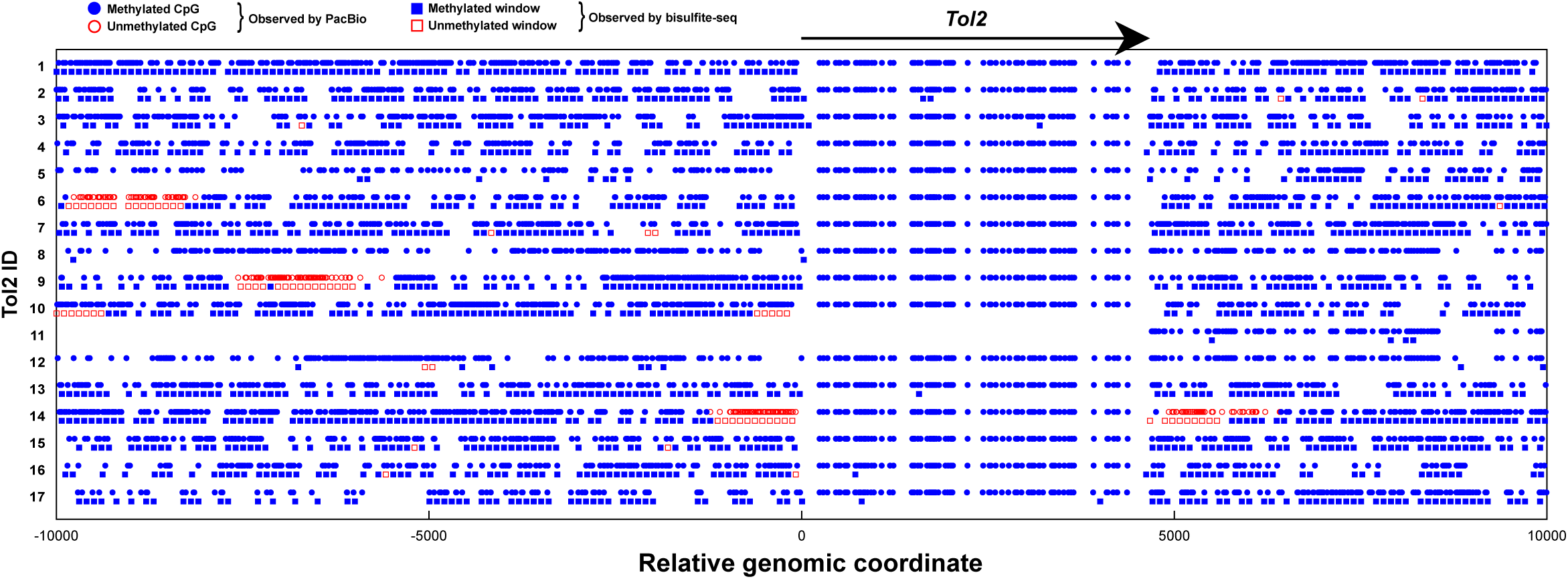
Methylation analysis of *Tol2*, a 4682-bp long autonomous transposon, in medaka. The new genome assembly of SMRT reads had 17 regions (contigs) that contained complete *Tol2* copies. The circles show our prediction of the methylation state of CpG sites, while the rectangles show the methylation states within each 100 bp window obtained from bisulfite sequencing. For both tracks, open/red indicates unmethylation and filled/blue indicates methylation. The arrow above indicates the region of *Tol2* insertions. As the eleventh region was located at the extreme of the contig, *Tol2* was not observed successfully by either SMRT sequencing or bisulfite sequencing. For the other 16 regions, methylation of *Tol2* was observed consistently by SMRT sequencing, while virtually no information was available on the *Tol2* region from bisulfite sequencing.

## 3 DISCUSSION

In this study, we addressed the problem of uncovering the landscape of DNA methylation of repetitive elements. To this end, we developed a unique application of SMRT sequencing to epigenetics. This direction had been already explored in the research community for bacterial and viral species. However, this application in large vertebrate genomes has been largely unexplored because of the subtle cytosine methylation signals in the kinetic information. Therefore, we proposed a new method to utilize relatively small amounts of kinetic information by incorporating a model reflecting our prior knowledge on the regional patterns of CpG methylation of vertebrate genomes. We confirmed the validity of our strategy by comparing the prediction to bisulfite sequencing data on medaka and to BeadChip analysis on human samples. These two datasets had very different characteristics, which seemed to be partly because of the methods used (*i.e.*, BeadChip was designed to observe mainly CpG islands that are often unmethylated, while bisulfite sequencing is used for genome-wide methylation analysis) and partly because of the nature of the samples used (*i.e.*, the medaka samples were derived from an inbred strain, while the human samples were from diploid cells). Despite such differences in characteristics, our method using the same parameters performed almost equally well for both datasets (Fig. 1F,G). These observations suggested that the choice of parameters is robust for a wide variety of samples, which is a desirable feature for any method. We also presented an extension of our method to accommodate intermediate methylation states, the discrete methylation level (DML), and confirmed a high correlation (*R* = 0.884) between DML and bisulfite methylation level.

We explored the epigenetic landscape of repetitive elements within the human genome. Using the hg19 reference genome is an apparent limitation. By assembling individual personal genomes instead of the reference genome, new insertions of these repetitive elements are expected to be found, and such active occurrences should be of interest. Indeed, we detected two novel LINE insertions that were estimated to be methylated. Importantly, the more recent the insertion event, the less divergent it would be from the original copy, and therefore, there would be less likelihood of it being anchored by short reads. In such cases, long SMRT reads shed new light on the ecosystem of active repetitive elements in personal human genomes.

Finally, our method had important strengths compared with conventional tools for epigenetic studies, such as bisulfite sequencing or affinity-based assays, with not only an expected increase in comprehensiveness by virtue of long SMRT reads, but also in the remarkable reduction of laboratory work. If an epigenetic study is conducted alongside a resequencing study or a *de novo* assembly study using SMRT sequencing, the methylation status could be called solely *in silico*, and no additional experiments would be necessary.

## METHODS

### Software availability

Our software program AgIn (<u>Ag</u>gregate on <u>In</u>tervals) is available at: <u>https://github.com/hacone/AgIn</u>

### Preparation of genomic DNA and SMRT sequencing

DNA was extracted from the testes of Hd-rR medaka with the DNeasy Blood & Tissue Kit (Qiagen, Hilden, Germany), following the tissue protocol. Genomic DNA was isolated from peripheral blood leukocytes of two Japanese patients using standard procedures after informed consent. The DNA featured A280/260 values of ∼1.8 and formed a clear, sharp band on agarose gel electrophoresis.

For the medaka sample and one human sample, genomic DNA was sheared using g-Tube devices (Covaris Inc., Woburn, MA, USA), targeting 20 kb fragments at 4300 rpm, 150 ng/μl and purified using 0.45*×* volume ratio of AMPure beads (Pacific Biosciences, Menlo Park, CA, USA). SMRTbell™ libraries were prepared with the DNA Template Preparation Kit 1.0 (Pacific Biosciences, Menlo Park, CA, USA) using the “20-kb Template Preparation using BluePippin Size Selection System (15 kb Size Cutoff)” protocol. Sequencing primer was annealed to the template at 0.833 nM concentration. SMRT bell™ templates were sequenced using magnetic bead loading, C4 chemistry, and polymerase version P6. Sequence data were collected on the magnetic bead collection protocol, 20 kb insert size, stage start, and 240 min movies in PacBio RS Remote.

For the other human sample, sequencing was performed as follows. Genomic DNA was sheared with using g-TUBE devices, targeting 10 kb fragments. SMRTbell™ libraries were prepared with the DNA Template Preparation Kit 2.0 (3∼10 kbp) (Pacific Biosciences, Menlo Park, CA, USA). Briefly, sheared DNA was end-repaired, and hairpin adapters were ligated using T4 DNA ligase. Incompletely formed SMRTbell™ templates were degraded using a combination of exonucleases III and VII. The resulting DNA templates were purified using (0.45*×*) SPRI magnetic beads (AMPure; Agencourt Bioscience, Beverly, MA, USA). Sequencing primers were annealed to the templates at a final concentration of 5 nM template DNA. SMRTbell™ library was sequenced using Magbead loading, C2 chemistry, and Polymerase version C2 or P4. Sequence data were collected on the PacBio RS for 120 min.

Regarding two human samples, the latter sample matches the one used for Illumina BeadChip analysis. We used the sequencing data and methylation state prediction from this sample solely for the analysis of intermediate methylation state prediction (Fig. 1G).

### Normalization of beta values of Illumina BeadChip

The respective beta values of an unmethylated CpG and a methylated CpG are not always equal to 0 and 1. Indeed, in our data, the distribution of raw beta values of Illumina BeadChip had bimodal peaks at 0.04 and 0.89. To compare beta values with our DML data (Fig. 1G), we treated 0.04 and 0.89 as unmethylated and methylated states respectively and normalized raw beta values by setting *x* ≤ 0.04 to 0, 0.89 ≤ *x* to 1, and 0.04 < *x* < 0.89 to (*x* − 0.04)/(0.89 − 0.04), proportionally.

### Raw IPDR and read coverage

We used the PacBio RS SMRT pipeline to process raw kinetic data from SMRT sequencing to obtain the mean IPDR and read coverage at each genomic position. *r*_*i*_ and 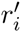 denote the mean IPDR associated with position *i* of the forward and reverse strands, respectively, and *c*_*i*_ and 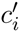 denote the read coverage at position *i* of the forward and reverse strands, respectively. To remove outlier noise inherent in raw data, mean IPDRs >10 were Winsorized to 10 and positions with less than three reads were excluded from the data (the latter was handled by SMRT Pipe). In bisulfite sequencing, CpG sites with ≥ 10 reads that mapped to C of either strand were considered covered. CpG sites that have a ≥ 1 position within a 21 bp window with ≥ 3 SMRT reads were counted as covered.

### Estimating the methylation status at individual CpG sites

Suppose that the focal genome has *n* CpG sites. We can assign identifiers ranging from 1 to *n* to individual *n* CpG sites and denote the genomic position of C of the *i*-th CpG site by *p*_*i*_. For example, the second CpG site at the 10th genomic position is denoted by “*p*_2_ = 10.” Our goal was to predict the methylation status, unmethylated or methylated, at *p*_*i*_ using information on read coverage and IPDR at positions surrounding *p*_*i*_. We used positions within 10 bases around *p*_*i*_ because these neighboring positions have proven to be effective in predicting 5-hydroxymethylcytosine, N4-methylcytosine, and N6-methylademine in bacteria genomes in previous studies (Flusberg et al., 2010; Clark et al., 2011, 2012; Fang et al., 2012). Neighboring positions are denoted by *p*_*i*_ + *j* for *j* = −10, …, +10 in the plus strand. For example, the positions 5 bases upstream and downstream of *p*_*i*_ are *p*_*i*_ − 5 and *p*_*i*_ + 5, respectively.

To achieve a better prediction, we derived a modified IPDR vector from raw read coverage and IPDR within 10 bases around *p*_*i*_. For this purpose, we took into account the property that any CpG site in one strand is reverse complementary to the CpG in the other strand, and the methylation status of Cs at a pair of CpG sites in both strands is consistent in most cases, making it meaningful to combine IPDR information for both strands to predict the methylation status. To represent positions in the minus strand, we note that since we set the position of C of the focal CpG in the plus strand to *p*_*i*_, the position of C of the CpG in the minus strand is *p*_*i*_ + 1, and the surrounding positions are *p*_*i*_ + 1 − *j* for *j* = −10, …, +10. In addition, we attached more importance to IPDR values associated with a higher read coverage and we quantified this as *c*_*p*_*i*_+*j*_ × *r*_*p*_*i*_+*j*_ in the plus strand (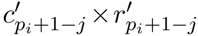 in the minus strand). We then took the sum of all the products and normalized it by dividing it by the total number of reads. Finally, we obtain the 21-dimensional modified IPDR vector for 21 genomic positions around CpG site *p*_*i*_:

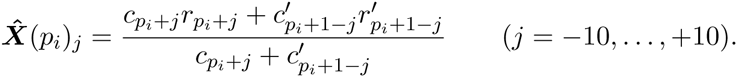

We are now in a position to define a classifier that uses 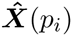 as explanatory variables and predicts the methylation status at *p*_*i*_, which is also estimated independently by bisulfite sequencing (Qu et al., 2012). We attempted to use linear discriminant analysis (LDA) with the discriminant function

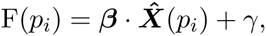

where we optimized values of the coefficient vector *β* and variable *γ* using bisulfite sequencing data as the training data set to improve the prediction. Supplemental Fig. S1A shows the optimized vector *β* that we used in this study. If the sign of the discriminant function, F(*p*_*i*_), is positive, the methylation status at *p*_*i*_ is defined as ‘methylated’; otherwise, it is defined as ‘unmethylated.’ We note that according to previous studies (Flusberg et al., 2010; Fang et al., 2012; Schadt et al., 2012), estimating 5-methylcytosine residues with low read coverage, for example 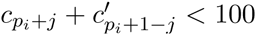 is prone to errors, demanding hundreds of reads, which is extremely costly to achieve.

### Predicting the methylation status of CpG blocks

In vertebrates, unmethylated CpG dinucleotides are rare (∼10%) and do not always exist in isolation, but they are likely to range over long unmethylated regions. This motivated us to integrate low-coverage reads around CpGs in a region to yield high-coverage for estimating the methylation status in the entire region, rather than at a single-base resolution. Let *A* denote a region. The following formula expresses the average IPDR vector for 21 genomic positions around all of the CpG sites in region *A* and its associated discriminant function:

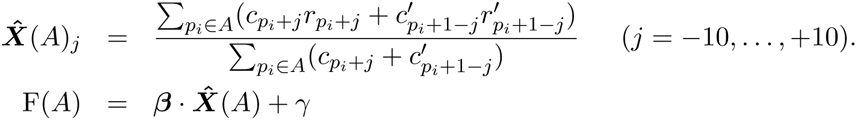

In processing a longer region with more CpG sites, the accuracy of methylation status prediction can improve, although smaller regions may be overlooked. In our analysis, we impose the constraint that each region has at least *b* CpG sites. For example, we can set *b* to 50 because Su *et al.* report that the average length of unmethylated regions in five human cell types is approximately 2 kb (Su et al., 2012b) and the average distance between neighboring CpG sites in the medaka genome is 53.5 bases, although this constraint should be adjusted according to each individual situation.

The possibility of the methylation (unmethylation, respectively) of *A* increases with a larger positive (negative) value of F(*A*), as well as for a larger total number of reads

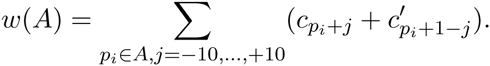

Thus, region *A* associated with a larger value of *w*(*A*)F(*A*) is better.

### Decomposing the genome into unmethylated/methylated CpG blocks

Now, we must consider how to decompose *n* CpG sites in the whole genome into methylated regions {*M*_*λ∈*Λ_} and unmethylated regions {*U*_*μ∈M*_} such that all regions are disjoint from each other, their union covers all CpG sites, and the two types of region occur alternatingly along the genome. To obtain better regions, we calculated the optimal decomposition of regions that maximizes the following objective function:

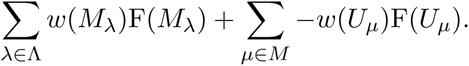

To solve this problem, we here mention one important characteristic of SMRT sequencing. Read coverage is not affected by the sequence composition in SMRT sequencing (Bashir et al., 2012; Zhang et al., 2012; English et al., 2012; Koren et al., 2012; Loomis et al., 2012). Thus, we assume that the sum of reads at the *j*-th position around all CpG sites in region *A* is a constant 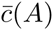 independent of *j*:

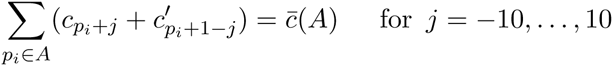

This allows us to transform *w*(*A*) into a simpler form:

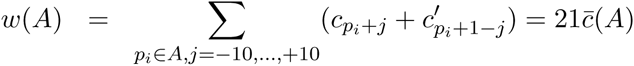

Subsequently, we can also simplify the objective function:

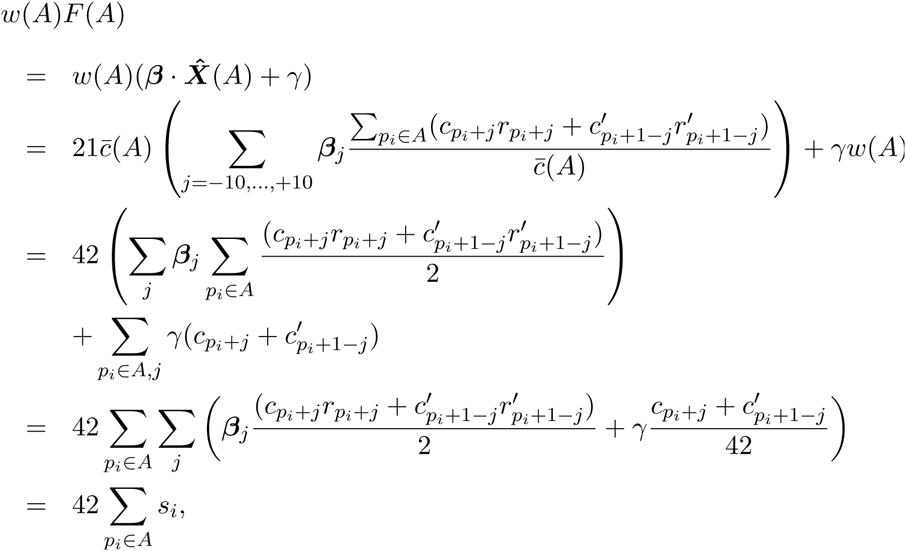

where *s*_*i*_ donates 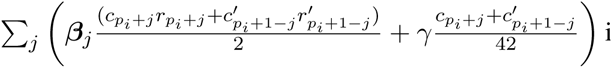 in the second last formula because the value only depends on read coverage and IPDR values at 21 genomic positions surrounding *p*_*i*_. Consequently, our objective function to optimize became a linear combination of *s*_*i*_:

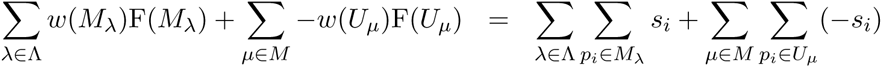

Although we set *s*_*i*_ to a score calculated from weighted IPDR information, we can set *s*_*i*_ to a log-likelihood function of the form - log *Q*_*i*_ for some likelihood function *Q*_*i*_.

This simple form motivated us to design a dynamic programming algorithm for calculating the optimal value efficiently. We considered the subproblem involving the first *i* CpG sites among all *n* sites, and let 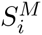 and 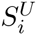 be the maximum value of the objective function when the last *i*-th CpG site was methylated and unmethylated, respectively. *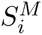* and 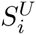 meet the following recurrences:

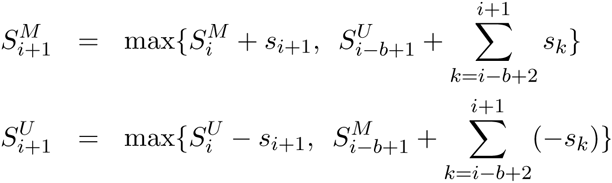

The first max term implies extension of the running region by one CpG site, while the second term means a switch from the previous methylation status and the initiation of a new region with ≥ *b* CpG sites. For example, we can set *b* to 50, but one can change the requirement for the minimum number of CpG sites in a region by making an appropriate adjustment to the second term. Of 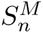 and 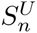, the larger value gives the maximum value, and tracing back the optimal path from the maximum value provides all the boundaries between neighboring methylated and unmethylated regions. To calculate regions satisfying the constraint on the minimum number of CpG sites, we generalized the dynamic programming idea proposed by Csűrös (Csűrös, 2004).

Thus far, we have assumed two possible methylation statuses, methylated or unmethylated, because this situation is true in most cases in our inbred strain sample (Qu et al., 2012). In human cells, however, many partially methylated cytosines have been reported (Lister et al., 2009). To consider such situations, we need to extend our algorithm to involve scores for three methylation statuses, methylated, unmethylated, and partially methylated. One can redesign the score function and the recurrence for each class. For example, making the parameters, *β*, *γ*, and *b*, depend on the class to which the *i*-th CpG site belongs, we can redefine the new recurrences for three classes:

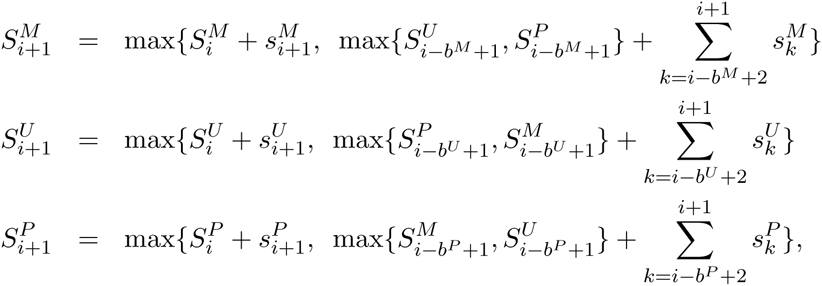

where *P* denotes “partially methylated,” *b*^*C*^ indicates the minimum region length for each class *C* ∈ {*M, U, P*}, and

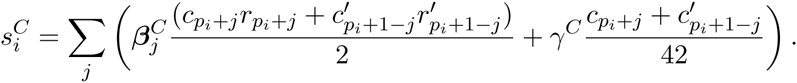

One might wonder if the hidden Markov Model (HMM) can be used for computing unmethylated and methylated regions; however, it is not obvious that using HMM guarantees the requirement that each range has ≥ *b* CpG sites.

### Computing discrete methylation levels

For calculating more quantitative methylation level named *discrete methylation levels*, we performed prediction using the set of 10 perturbed intercept values (*γ* ranging from −12% to +24% by 4%) so we obtain 10 predictions on each CpG site. Then, on each CpG site, the number of predictions that favored methylation were divided by 10, yielding the discrete methylation level ranging over [0, 1].

### Methylation status calculated from bisulfite sequencing

We evaluated the prediction accuracy of our integration method using methylation scores calculated from bisulfite-treated Illumina reads as the answer set. Let *S* be the set of bisulfite-treated Illumina reads covering the *i*-th CpG site, *x* be the number of methylated CpGs in *S* at *i*, and *y* be the coverage of *S* at *i* (the size of *S*). We defined the methylation score at *i* as *x/y*. We then defined the methylation status as ‘unmethylated’ if the score was less than 0.5; otherwise, it was defined as ‘methylated’.

We need to carefully constrain the value of the coverage *y*. Allowing a lower value of *y* is likely to produce more erroneous methylation scores, while using *y* greater than a higher threshold would reduce the number of CpGs associated with their methylation scores. The average coverage was 9.40-fold in our bisulfite-treated reads collected from testes of the Hd-rR medaka inbred strain; however, the coverage at individual CpG sites varied to some extent. We defined the methylation score only when the CpG site was covered by 10 or more reads (*i.e.*, *y ≥* 10) so that make sure the scores are robust enough.

### Prediction accuracy of our method at individual CpG sites

We predicted the methylation status of each CpG site by checking whether the CpG site was located in an unmethylated or methylated region according to the output of our integration method. We measured the accuracy of the prediction by checking the consistency between the prediction and the methylation score associated with each CpG site. CpG sites without methylation score (due to the lack of bisulfite-treated reads) were ignored. We treat a unmethylated status as positive and a methylated status as negative, because we are more interested in identifying rare unmethylated regions accounting for a small portion (e.g., ∼10%) of CpG sites.

### Methylation analysis of human repetitive elements

We started the analysis by listing repetitive elements using the Repeat Library “20140131” (Smit, A., Hubley, R. & Green, P. Repeatmasker open-4.0 at http://www.repeatmasker.org). Only repetitive elements containing at least 10 CpG sites were considered. We calculated the methylation levels of CpG sites as discrete methylation levels (DML), and CpG sites with a DML<0.4 were considered as unmethylated. To further reduce the degree of unmethylation assigned false-positively, we filtered out repetitive elements with an average read coverage on CpG sites of <5.0. Finally, we treated repetitive elements as unmethylated if more than half of the CpG sites were unmethylated; otherwise, they were considered as methylated.

The relationships between methylation state and CpG density or divergence were tested for statistical significance using the Mann-Whitney U test. To draw ideograms in Figure 2C, we counted the numbers of unmethylated and methylated repeats in every 5 Mb bin and then used the Ideographica web server (Kin and Ono, 2007) to generate the images. In Supplemental Figure S4, Kernel PCA analysis was performed using spectrum kernel. As the magnitude of sequence divergence among occurrences was markedly variable for different types of repetitive elements, it was necessary to optimize the k-mer size for each type of repetitive element to achieve better visualization.

### Validation of our prediction by bisulfite Sanger sequencing

Bisulfite conversion of genomic DNA was performed using a commercially available kit (MethlEasy Xceed Rapid DNA Bisulphite Modification Kit; Human Genetic Signatures, NSW, Australia). Briefly, 5 μg of DNA was denatured by 0.3 M NaOH for 15 minutes at 37*°*C. Subsequently, the samples were incubated with bisulfite solution for 45 minutes at 80*°*C. After purification, the eluted samples were incubated for 20 minutes at 95*°*C. The converted DNA was stored at −20*°*C for PCR amplification.

To perform targeted PCR on the 21 regions selected for validation, we designed primers for nested PCR to amplify 111∼622bp fragments of bisulfite-converted DNA (Supplemental Table S2). Primer pairs were purchased from Life Technologies (Supplementary Information). PCR was performed in a volume of 50 μL containing 1 *×* EpiTaq PCR Buffer, 2.5 mM MgCl2, 0.3 mM dNTP mix, 20 pmol primers, 1.25 units TakaraEpiTaq HS polymerase (Shiga, Japan), and 50 ng bisulfite-converted DNA. PCR conditions were 40 cycles of 98*°*C for 10 seconds, 55*°*C for 30 seconds, and 72*°*C for 1 minute. To check the quality of the PCR products, 2% agarose gel electrophoresis was used in 1 *×* TAE buffer at 50 volts for 15 minutes. The amplified products were visualized using a LED transilluminator, and the product bands were purified using the NucleoSpin Gel and PCR Clean-up kit (Macherey-Nagel GmbH & Co. KG, Dueren, Germany). Targeted PCR products were sequenced directly using ABI3730 sequencers with BigDye v3.1 chemistry (Applied Biosystems, Foster City, CA, USA).

Finally, we processed the obtained sequencing data using the QUMA online tool (Kumaki et al., 2008) for analysis and visualization of the methylation patterns (Supplemental Fig. S3).

### Methylation analysis of medaka *Tol2* elements

In Figure 3, we applied our method to observe the methylation state of a new medaka assembly. For comparison, we also called the methylation state on every 100-bp window using Bismark software and the publicly available bisulfite-treated reads from the testes of the Hd-rR strain. Among the assembly, we identified 17 contigs containing *Tol2* elements by BLAST search.

### Other data sources and data visualization

Figure 1B and Supplemental Figures S5-S6 were produced using the UCSC Genome Browser (http://genome.ucsc.edu/) (Karolchik et al., 2014). We used human bisulfite sequencing data and unmethylated regions available in the GEO database (Song et al., 2013).

## DATA ACCESS

The sequence data (SMRT reads) from the medaka sample are deposited at the NCBI Archive (Accession No. SRP020483). Sequence data from a Japanese individual are available under controlled access through the National Bioscience Database Center (NBDC, accession number JGAS00000000003).

## ACKNOWLEDGMENTS

This work was supported in part by MEXT (Grants-in-Aid for Scientific Research 22129008 and 23241058 [SM]). We would like to thank Ryohei Nakamura for stimulating discussions.

## Legends for supplemental tables and figures

**Supplemental Table S1.**
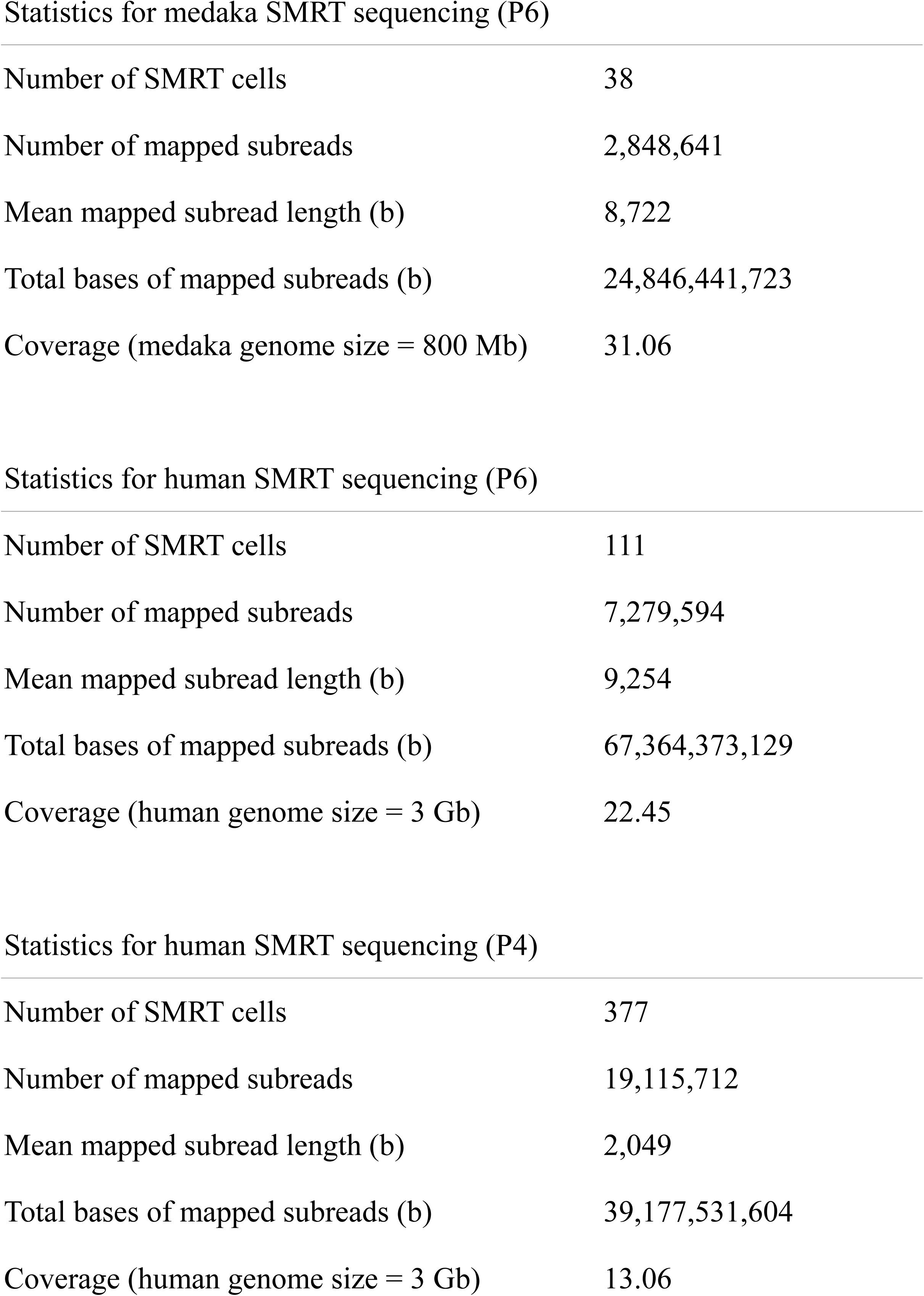
Statistics of SMRT sequencing data production. Summary statistics of SMRT sequencing data collected in this study.

|  |  |
| --- | --- |
| Number of SMRT cells | 38 |
| Number of mapped subreads | 2,848,641 |
| Mean mapped subread length (b) | 8,722 |
| Total bases of mapped subreads (b) | 24,846,441,723 |
| Coverage (medaka genome size = 800 Mb) | 31.06 |
| Statistics for human SMRT sequencing (P6) |  |
| Number of SMRT cells | 111 |
| Number of mapped subreads | 7,279,594 |
| Mean mapped subread length (b) | 9,254 |
| Total bases of mapped subreads (b) | 67,364,373,129 |
| Coverage (human genome size = 3 Gb) | 22.45 |
| Statistics for human SMRT sequencing (P4) |  |
| Number of SMRT cells | 377 |
| Number of mapped subreads | 19,115,712 |
| Mean mapped subread length (b) | 2,049 |
| Total bases of mapped subreads (b) | 39,177,531,604 |
| Coverage (human genome size = 3 Gb) | 13.06 |

**Supplemental Table S2.**
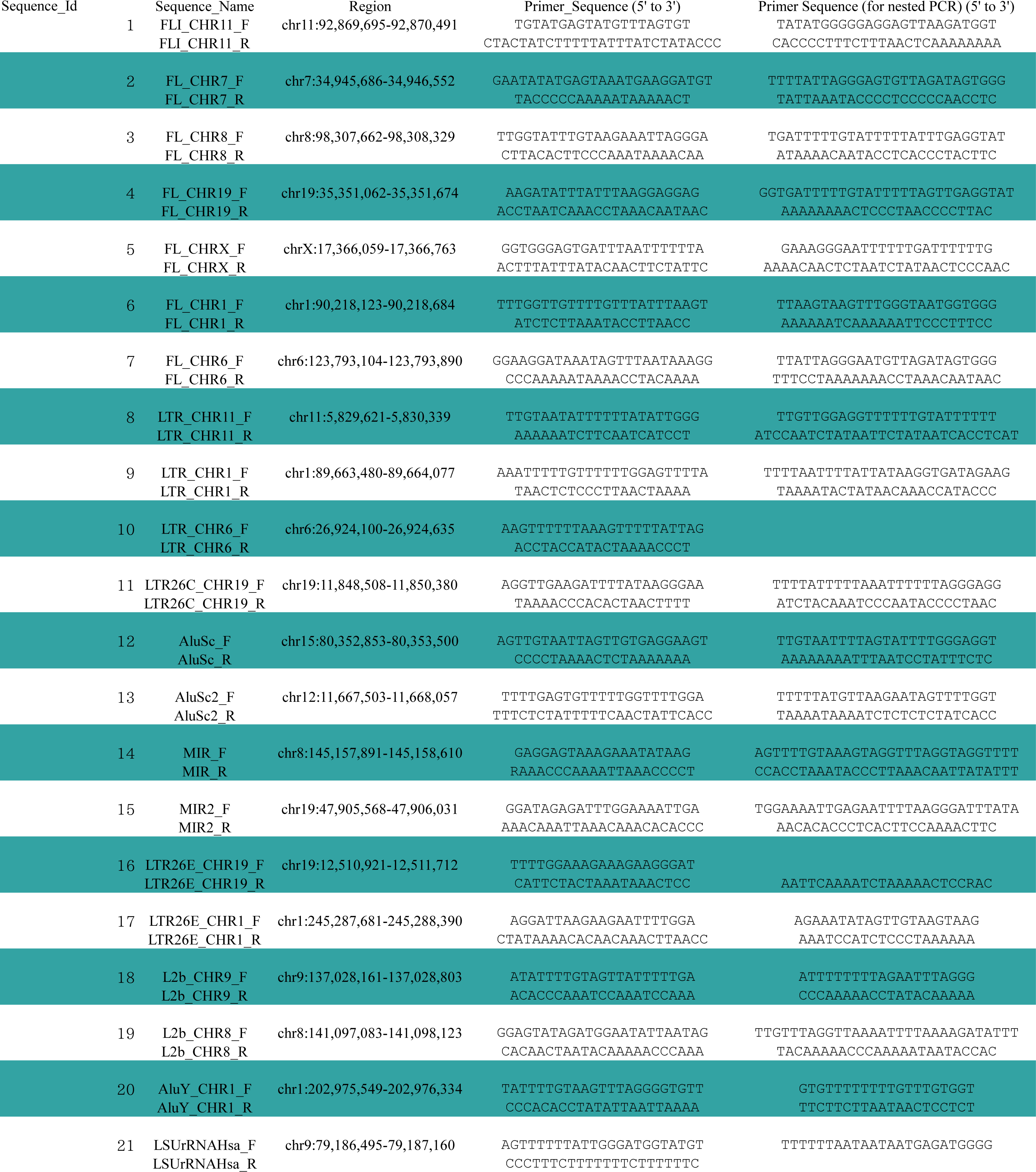
The primers for nested PCR of the bisulfite treated blood DNA. The primers for nested PCR are shown alongside the sequence IDs that correspond to those in Supplemental Figure S4, the sequence names, and the target genomic regions. For each entry, the forward primers appear in the top row, and the reverse primers appear in the second row.

**Supplemental Table S3.** DNA methylation states of full-length LINE/L1 elements. According to the three classes of full-length LINE/L1 elements in L1Base, we examined DNA methylation states of LINE/L1 elements in each class.

| Class | With >9<br>CpGs (A) | Covered<br>with SMRT<br>reads (B) | B/A | Covered<br>with >5x<br>(C) | C/A | Unmethylated<br>(D) | D/C |
| --- | --- | --- | --- | --- | --- | --- | --- |
| non-intact L1s due to multiple mutations | 11646 | 11497 | 98.72% | 10465 | 89.86% | 64 | 0.61% |
| L1s with an intact ORF2 but a disrupted ORF1 | 103 | 100 | 97.09% | 91 | 88.35% | 0 | 0.00% |
| L1s with intact in the two ORFs | 145 | 133 | 91.72% | 119 | 82.07% | 0 | 0.00% |

**Supplemental Figure S1.**
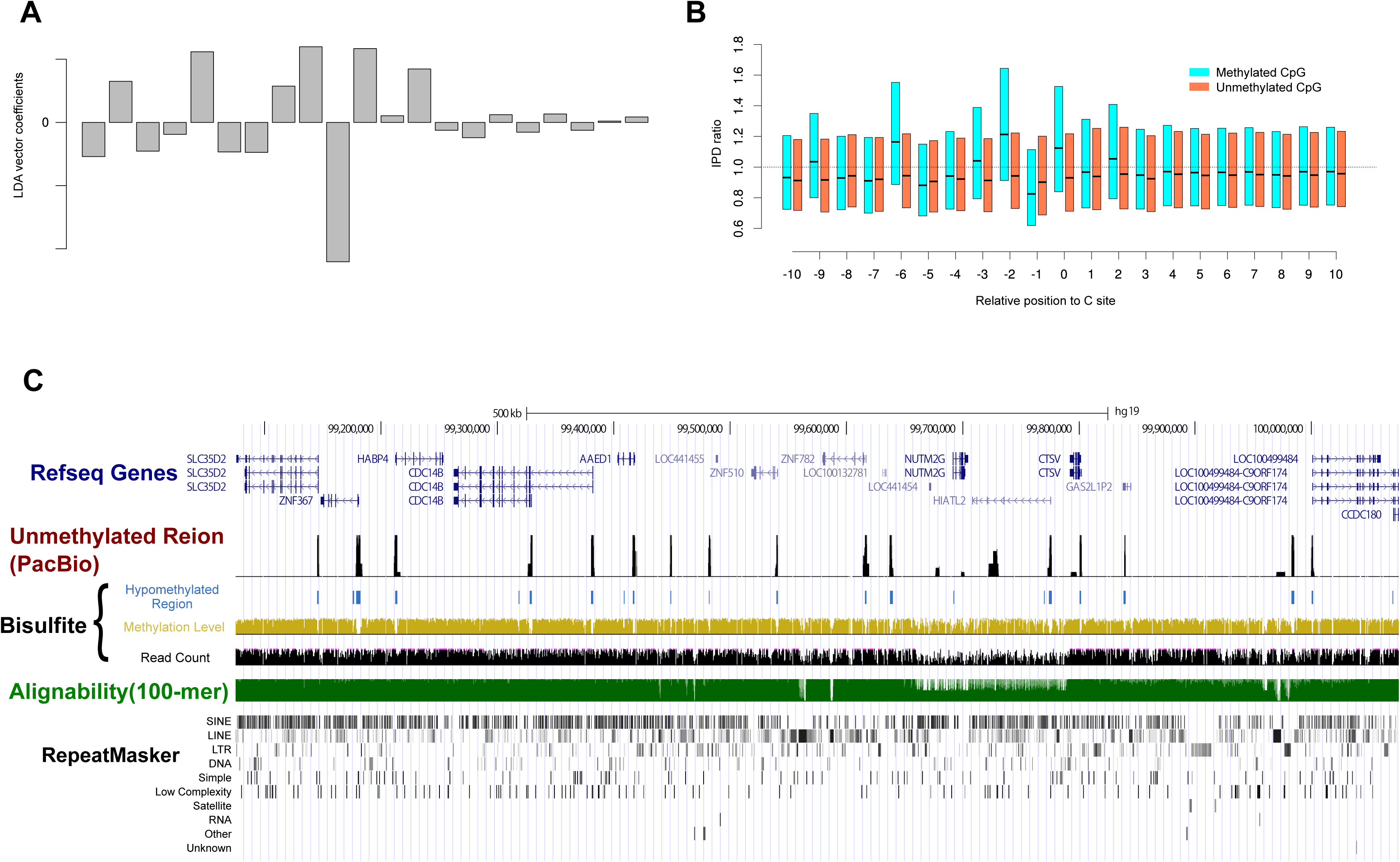
The normal vector used for prediction. **A.** The normal vector *β* used for prediction. We calculated *β* as follows. Firstly, we classified the CpGs on the scaffold 1 in the medaka Hd-rR genome (version 1) into methylated CpGs and unmethylated CpGs according to bisulfite sequencing data. Next, for each CpG site, we calculate the IPD ratio profiles as the 21-dimensional vectors based on SMRT sequencing kinetics data. Then, using LDA (Linear Discriminant Analysis), we tried to find the best hyperplane that could separate these IPD ratio profiles into each class, namely, methylated or unmethylated. The normal vector of this hyperplane is denoted by *β*. **B.** The average IPDR profiles around unmethylated and methylated CpG sites. The x-axis shows the positions within 10 bp of the focal CpG site at the position represented by 0. The y-axis indicates IPDR values. The red- and blue-colored box plots at each position show the distributions of IPDR values around unmethylated and methylated CpG sites, respectively. The bottom, middle and top of each box plot indicate the first, second, and third quartiles, respectively, of the distribution. **C.** An example in which both our method and bisulfite sequencing are consistent in showing unmethylation in gene promoters. The tracks are similar to those in Figure 1B.

**Supplemental Figure S2.**
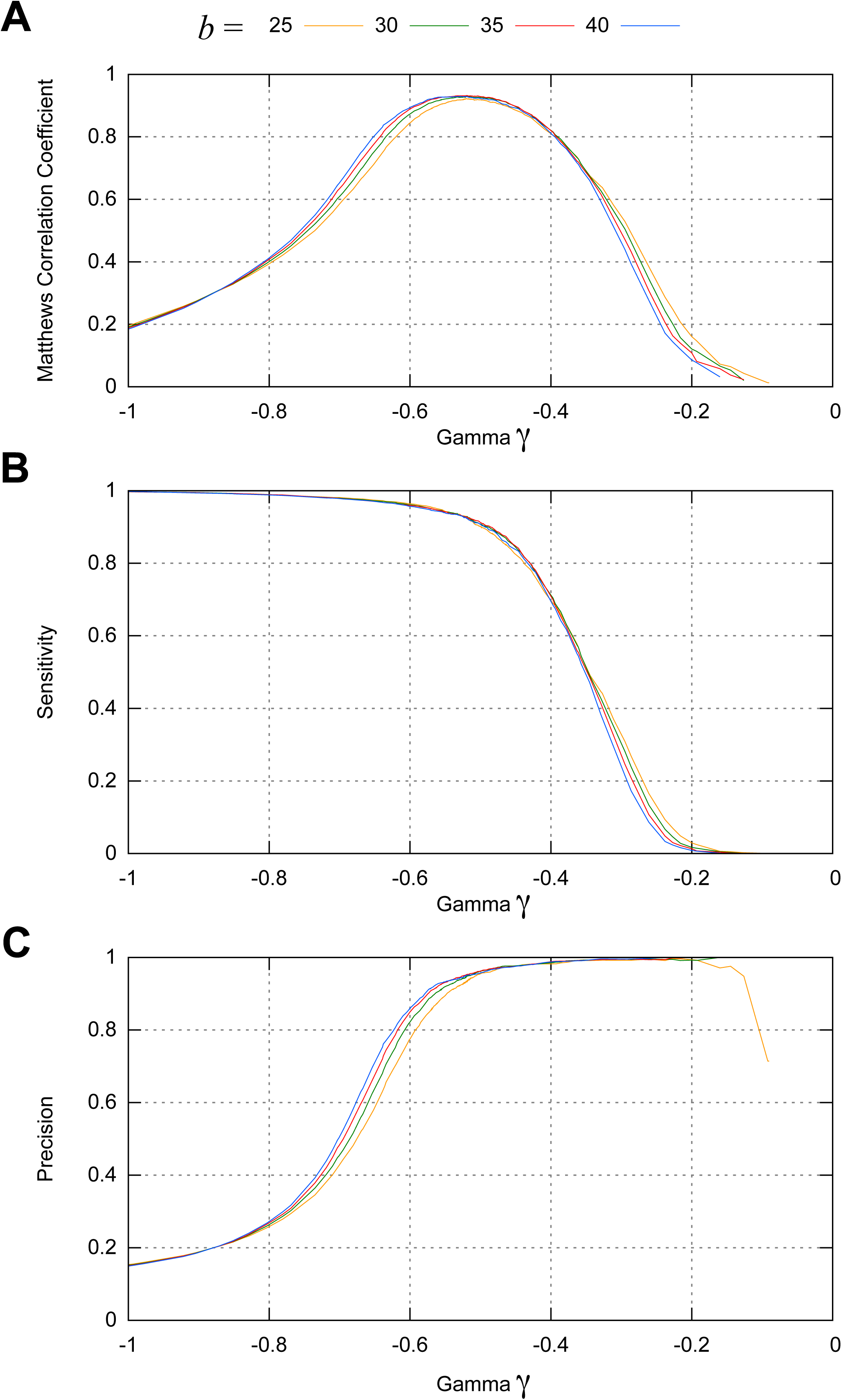
Accuracy measures on the chromosome 1 of the medaka Hd-rR genome (version 2) **A-C.** Matthew’s correlation coefficient **(A)**, sensitivity **(B)**, and precision **(C)** as a function of the intercept of the hyperplane *γ*, on the chromosome 1 in the medaka genome (version 2) with a 29.9-fold mapped read coverage. Matthew’s correlation coefficient represents an overall accuracy of our prediction. The differently colored curves correspond to the different lower bound of number of CpG sites, denoted by *b*, that was used for the prediction. Our prediction achieved 93.0% sensitivity and 94.9% precision at *b* = 35 and *γ* = −0.526. Or sensitivity (93.67%) and precision (93.88%) are close to each other when *b* = 35 and *γ* = −0.540.

**Supplemental Figure S3.**
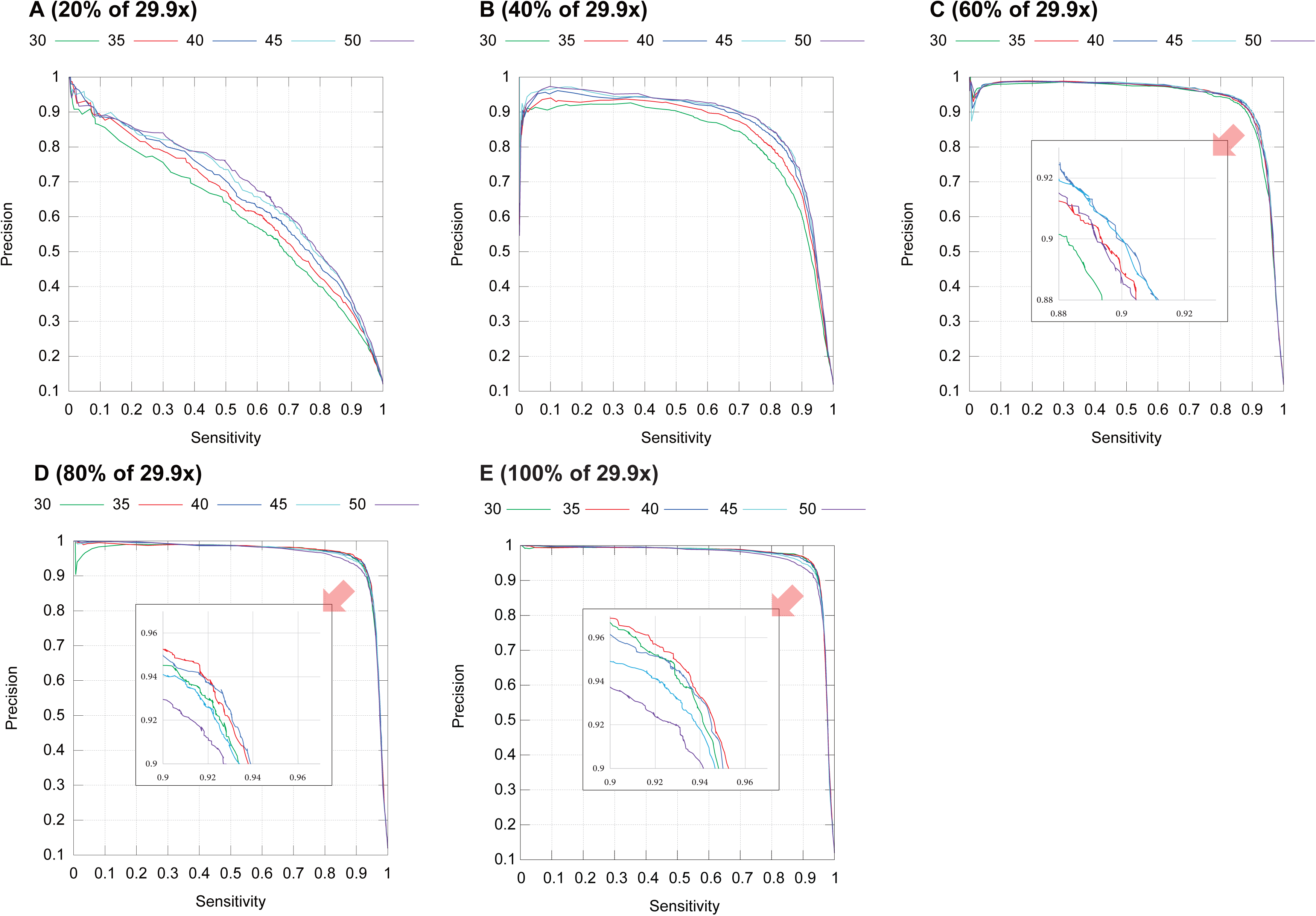
Sensitivity and precision of predicting unmethylated regions with ≥ *b* CpG sites for a variety of read coverages. We continue to use *b* to denote a lower bound of the number of CpG sites in a region. For *b* = 30, 35, 40, 45, 50, we plot the sensitivity and precision curves when the read coverage is 20% of 29.9x (**A**), 40% of 29.9x (**B**), 60% of 29.9x (**C**), 80% of 29.9x (**D**), and 29.9x (**E**). The sensitivity and precision were evaluated on the chromosome 1 of the medaka Hd-rR genome (version 2). For better prediction with a smaller coverage, a wider window was favored. Precisely, setting *b* to 50 outperforms the other values for coverages, 20% and 40%, but it becomes inferior for 80% and 100%. In contrast, both sensitivity and precision increase for larger coverages, 80% and 100%, when *b* is set to smaller values, 35 and 40. In particular, Figure **E** shows that for coverage 100% (29.9x), setting *b* to 35 is better than other values of *b*. Figure **C** also highlights that even with a small coverage 60% of 29.9x, both sensitivity and precision are ∼ 90% for *b* = 45.

**Supplemental Figure S4.**
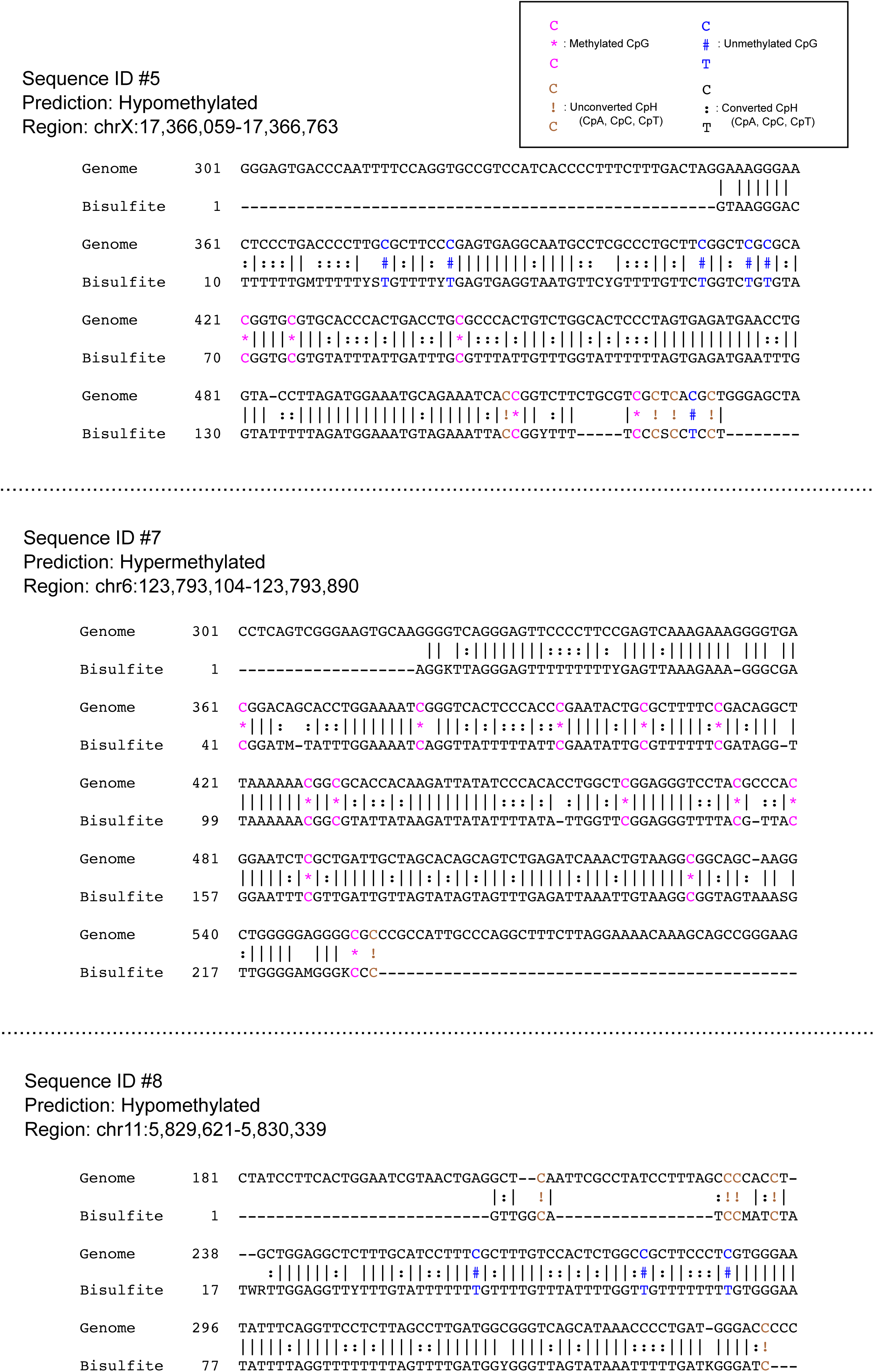
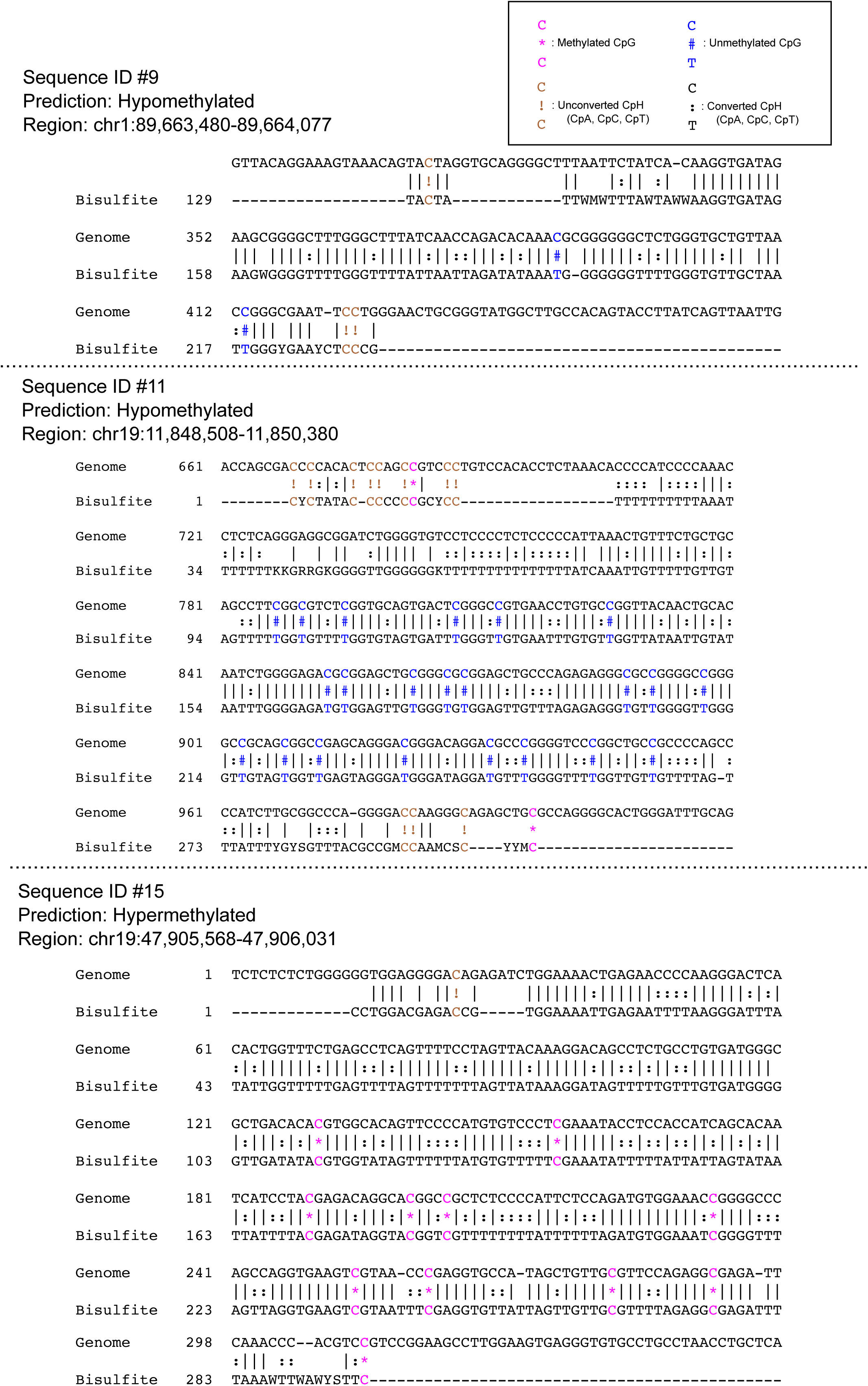
Methylation analysis of selected regions for validation of our prediction. Of the 21 regions selected for validation of our method, 6 were amplified, and their Sanger sequencing reads were aligned to the target regions. In the alignments, the methylated (unconverted) CpGs are represented by the pink asterisks (*), and the unmethylated (converted) CpGs by the blue number sign (#). We can assess the efficiency of bisulfite conversion and the quality of the alignment by looking at non-CpG C sites (CpHs) because Cs in CpHs are usually unmethylated and should always be converted to Ts (represented by the colons (:)). Thus unconverted CpHs, which are highlighted by the brown exclamation marks (!), indicate low quality regions. The solid lines represent the other types of matches.

**Supplemental Figure S5.**
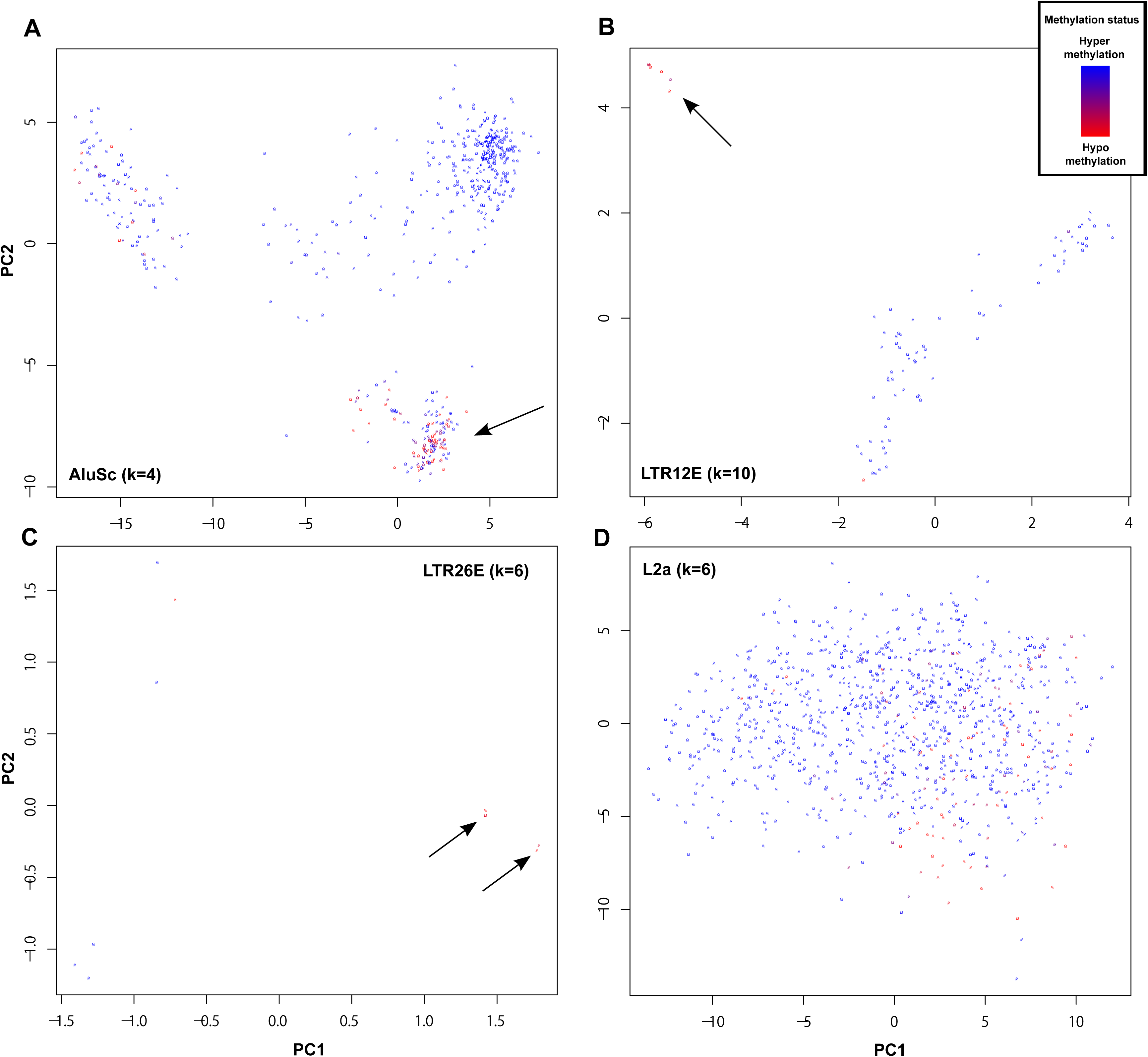
Kernel PCA analysis of sequence feature and methylation state. The results of Kernel PCA analysis are shown for 4 selected classes of repetitive elements, AluSc **(A)**, LTR12E **(B)**, LTR26E **(C)**, and L2a **(D)**. We projected the repeat occurrences into the plane based on the distance metrics that we defined using the spectrum kernels and their top 2 principal components. The colors of the dots represent the methylation state of the repeat occurrences; namely, red indicates unmethylation and blue methylation. The arrows show the unmethylated occurrences that are clustered in terms of the sequence features.

**Supplemental Figure S6.**
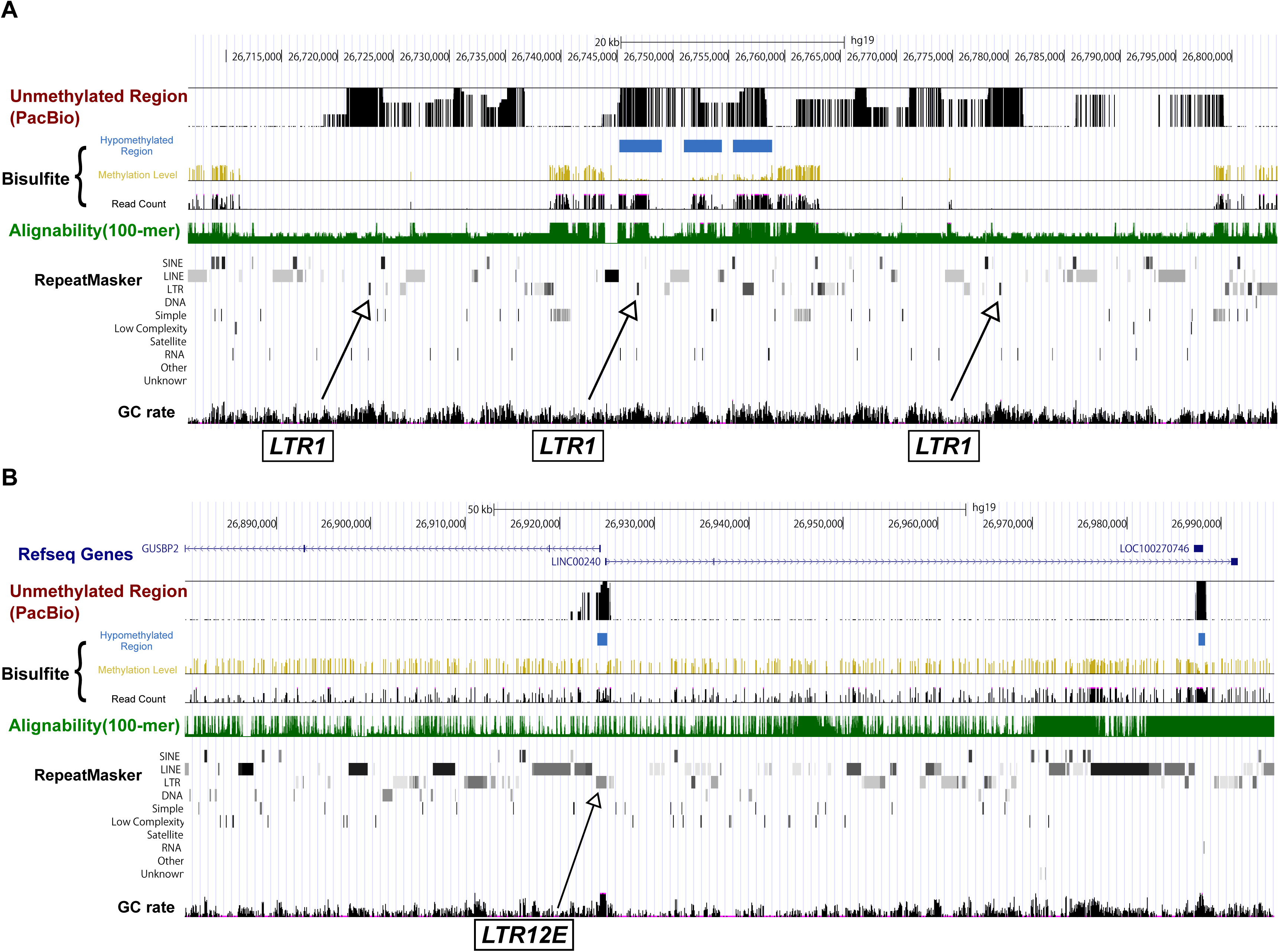
Examples of unmethylated repeat occurrences in a unmethylation ‘hot spot’. Three adjacent LTR1 elements were unmethylated in this region **(A)**, and a LTR12E element was located at a unmethylated bi-directional promoter region **(B)**. Both regions are on the p-arm of the chromosome 6. The arrows indicate the locations of LTR1 and LTR12E. From top to bottom, below the RefSeq gene track, black bars indicate unmethylated regions predicted from SMRT sequencing data using our method. Yellow and black bars show the methylation level and read coverage obtained from public bisulfite sequencing data, respectively, and blue boxes show unmethylated regions predicted from the bisulfite data. Green bars below indicate the alignability of short (100-bp) reads. The bottom rows shows repeat masker tracks and GC rate for every 5 bp window.

**Supplemental Figure S7.**
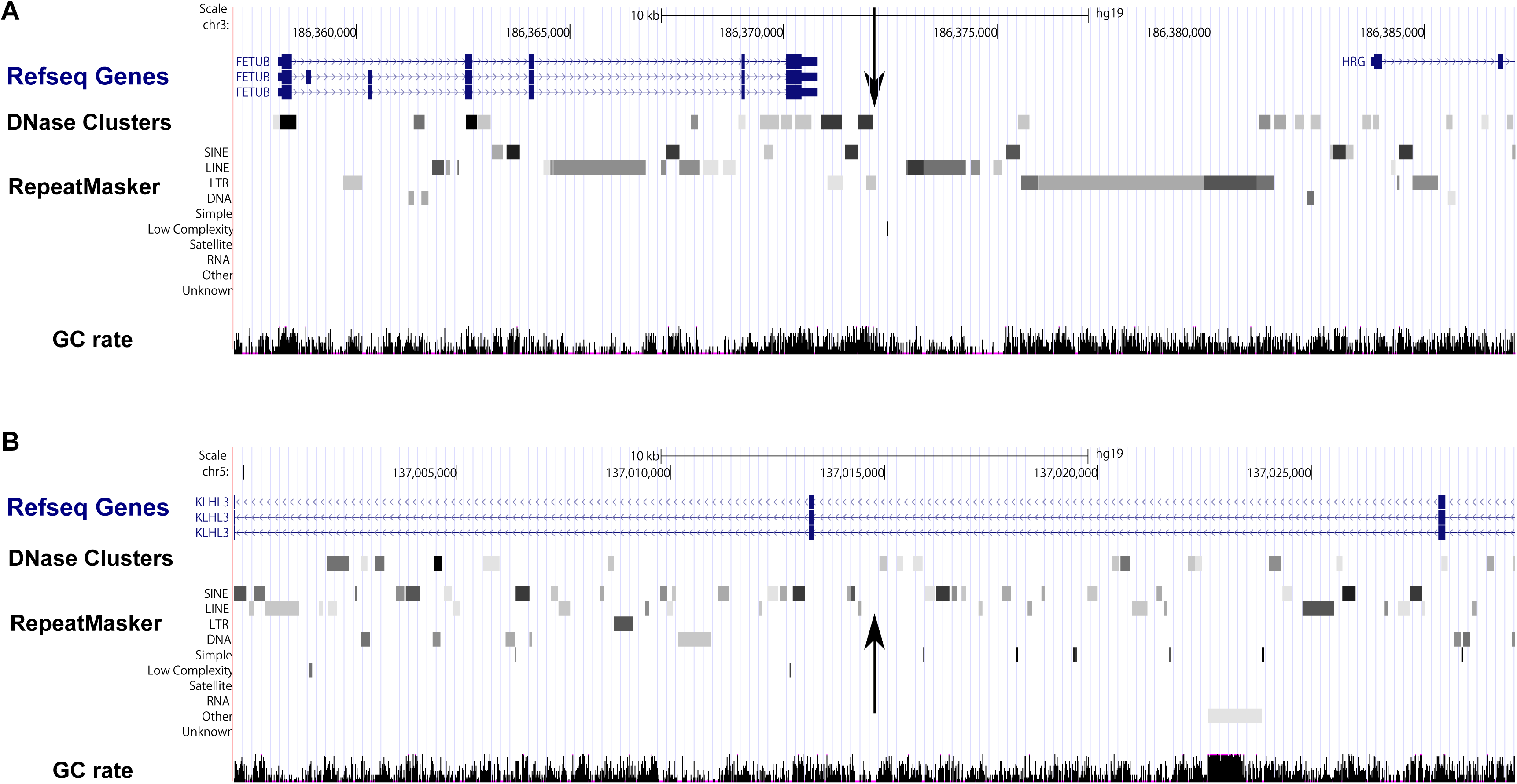
Two LINE insertions novel to hg19. We identified two LINE insertions by comparing a new assembly obtained from SMRT reads and the hg19 reference genome. The vertical arrows indicate the locations of the identified novel insertions. Specifically, one is aligned at 186,372,132 in Chromosome 3 with identity 99.02%, and the other at 137,014,775 bp in Chromosome 5 with identity 98.71%. From top to bottom, the tracks shown are RefSeq genes, DNase clusters, repeat masker masked regions, and GC rate for every 5 bp window.

